# Invasion by *Fusobacterium nucleatum* Causes Upregulation of *dll4* and *klf4* Expression in Caco2 Cells

**DOI:** 10.1101/2020.10.26.354910

**Authors:** Kyla Cochrane, Avery V. Robinson, Jacqueline Powers, Scott D. Brown, Robert A. Holt, Emma Allen-Vercoe

**Affiliations:** Canada’s Michael Smith Genome Sciences Centre, BC Cancer, Vancouver, British Columbia, Canada; Department of Molecular and Cellular Biology, University of Guelph, 50 Stone Road East, Guelph, Ontario Canada; Department of Medical Genetics, University of British Columbia, Vancouver, British Columbia, Canada; Department of Molecular Biology & Biochemistry, Simon Fraser University, Burnaby, British Columbia, Canada

**Keywords:** *Fusobacterium nucleatum*, colorectal cancer, transcriptome, host cell infection

## Abstract

*Fusobacterium nucleatum* is an emerging microbe of importance in the pathogenesis of colorectal cancer. Strains of this enigmatic bacterial species vary in their capacity to invade human epithelial cells, a virulence determinant which has important implications in disease. Here, we infected human colorectal epithelial (Caco-2) cells *in vitro* with a known, highly invasive strain of *F. nucleatum* isolated from a Crohn’s Disease patient, as well as a further invasive isolate of *F. nucleatum* derived from a colorectal cancer tumour. We used transcriptional profiling to determine the human genes upregulated during the invasion process compared to exposure to a non-invasive *E.coli* control strain. Infection with *F. nucleatum* strains resulted in the upregulation of several host genes, including two associated with tumorigenesis: *dll4* and *klf4*.

## Introduction

*Fusobacterium* is a heterogeneous, Gram-negative, anaerobic, bacterial genus with varying degrees of virulence observed among isolates. *Fusobacterium nucleatum* is found in the gut microbiome as well as other niches in the body. Virulent isolates have been implicated in colorectal cancer (CRC) development and are associated with poor prognosis (reviewed by Gethings-Behncke *et al*.(1)). Found in primary colorectal carcinoma tissues, *Fusobacterium* is detectable across disease stages (reviewed by Brennan and Garrett(2)). Intracellular *Fusobacterium* increases tumor cell proliferation *in vitro* and *in vivo* (reviewed by Ternes *et al.*(3)). It is hypothesized that the invasion of virulent *Fusobacterium* into the host cells alters the gene expression in both bacteria and host, setting into motion, or otherwise potentiating, a chain of events that can lead to CRC.

Our previous transcriptomic study assessed changes in *F. nucleatum* gene expression co-incident with host cell invasion in an effort to identify potential bacterial virulence factors associated with invasion (4). Here we use the same transcriptomic approach to investigate changes in human gut epithelial cell gene expression upon invasion with CRC associated *F. nucleatum.* The results of our comparative whole transcriptomic (RNA-seq) analysis using Caco-2 cells after exposure to *F. nucleatum* isolates provide insight into host cell expression changes that occur directly after invasion.

## Methods

Bacterial culture: *Fn* subsp. *animalis* 7-1 (*Fn*7-1) 7-33C1 (*Fn* 7-3) were used in these experiments, as well as *Escherichia coli* strain Nissle 1917. *Fn7-1* has been previously isolated, extensively phenotyped and sequenced by our group (5, 6). Although not specifically CRC-associated, *Fn* 7-1 has been extensively profiled in terms of adhesion and invasion assays (6) and has recently been shown to induce tumorigenesis in C57BL/6J–*Apc^Min/+^* mice (7). *Fn*7-3 was isolated directly from a CRC tumor biopsy through culture on selective medium (4). *E. coli* K-12 (Sigma-Aldrich) was used as a control strain. Propagation of bacterial strains was carried out in tryptic soy broth supplemented with hemin (5μg/mL) and menadione (1μg/mL) (TSB_supp_) (Sigma-Aldrich), with incubation in a humidified anaerobic chamber (Ruskinn Bug Box) at 37°C under an atmosphere of N_2_:CO_2_:H_2_ 90:5:5. For infection assays, culture grown to late log phase and normalized for cell number using McFarland standards.

Caco-2 Cell culture:Caco-2 human colon adenocarcinoma cells (American Type Culture Collection (ATCC), Manassas, VA line HTB37™) were cultured in Dulbecco’s Modified Eagle Media (DMEM) (Sigma-Aldrich) supplemented with 10% fetal bovine serum (FBS) (ThermoFisher), 10mM sodium pyruvate (Sigma-Aldrich), and 5μg/mL Plasmocin (Invitrogen) at 37°C in 5% CO_2_. For consistency, only Caco-2 cells from passages 4-12 were used for experiments.

Infection assay: Caco-2 cells were grown to 85% confluence, washed briefly, infected to a multiplicity of infection of 100:1 with bacterial cells and incubated at 37°C in 5% CO_2_ for 4 hours (6). Cells were then washed and treated with fresh DMEM containing gentamicin (0.5mg/mL) (Sigma-Aldrich) for 30 minutes at 37°C in 5% CO_2_, then trypsinized and quenched. All subsequent steps were carried out using reagents at 4°C. RNA was stabilized through the addition of TRIzol® Reagent (1mL per 50-100mg of pelleted sample) (ThermoFisher), before storage at −80°C. RNAlater^®^ (Qiagen) was used in all buffers throughout the extraction process. Total RNA extraction was performed within 15 minutes to ensure minimal transcriptional changes during the process. Three biological replicates were performed.

Transcriptional profiling: Strand-specific RNA-seq libraries were constructed as previously described (4), and sequenced on the Illumina HiSeq 2000 platform, yielding an average of 51.4 million reads per replicate. After filtering and alignment, host cell gene expression values (RPKM) were determined and differential gene expression (fold-change) between *F. nucleatum* infected or *E. coli* K12 control and non-infected host cells was calculated. Differentially expressed genes (DEG) were defined as those that met the arbitrary thresholds of RPKM ≥0.1, a minimum of 10 mapped sequence reads, an uncorrected p-value < 0.05 and an absolute fold-change value of at least four. Guided by the results of differential expression analyses, we selected a subset of 15 cancer relevant DEGs to investigate further. We designed a custom CodeSet® for the NanoString® nCounter assay to verify RNA-seq derived differential gene expression values. Next, we used the Caco-2 invasion assay and NanoString® profiling to explore modulation of these genes by different strains of *F. nucleatum* and the *E. coli K12* control. Further analysis using qPCR was performed on human genes of interest.

## Results

We detected 58 up-regulated genes and 12 down-regulated genes by these criteria (Figure 1; Panel A). Using NanoString® to hone in on 15 cancer-related genes, we show expression differences are highly concordant with RPKM values from RNA-seq (Pearson’s correlation coefficient R=0.97, p=2.2×10^−9^; Figure 2). Leveraging our custom CodeSet® across microbial strains allowed us to identify two genes, *dll4* and *klf4*, that were up-regulated in Caco-2 cells incubated with both *F. nucleatum* isolates relative to the lab-adapted *E. coli* K12 isolate (Figure 1; Panel B). qPCR further resolved that while the upregulation of *dll4* is significantly greater than that of Caco-2 cells incubated with *E. coli* K12 (p<0.0001), the upregulation of *klf4* is more limited (Figure 1; Panel C).

**Figure 1.**
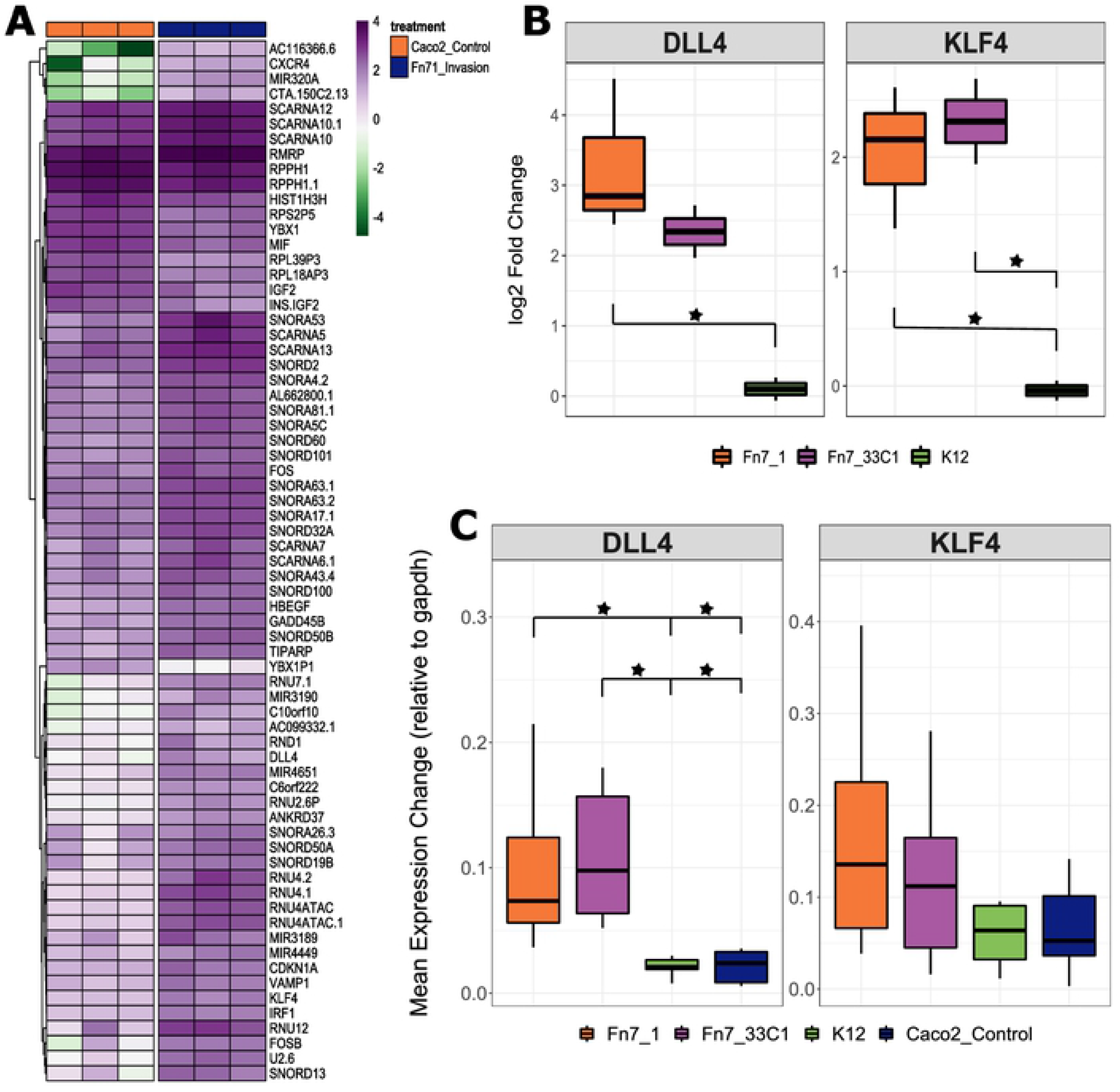
**Panel A:** Heatmap highlighting 58 up-regulated and 12 down-regulated host genes during Fusobacterial infection as determine by RNA-seq. Data is shown as log_2_RPKM. **Panel B:** Boxplot depicting the two human genes of interest, determined using a log fold change approach, our custom designed NanoString CodeSet® and *Fusobacterium*/*Escherichia* strain infection assays of Caco-2 cells. A statistically significant difference was observed using ANOVA and Tukey Post-Hoc analysis between Fn7-1 and *E. coli* K12 for KLF4 (p = 0.0249) and DLL4 (p=0.0298). For the comparison between Fn7-3 and *E. coli* K12, only KLF4 showed a significant difference (p= 0.0226). **Panel C:** For further resolution and to increase the number of replicates, qPCR TaqMan assays were used to further validate our two genes of interest: *dll4* and *klf4*. A statistically significant difference was observed using ANOVA and Tukey Post-Hoc Analysis between Fn7-1 or Fn-7-3 and either *E.coli* K12 or uninfected Caco2 cells for DLL4 only (p<0.0001).

**Figure 2.**
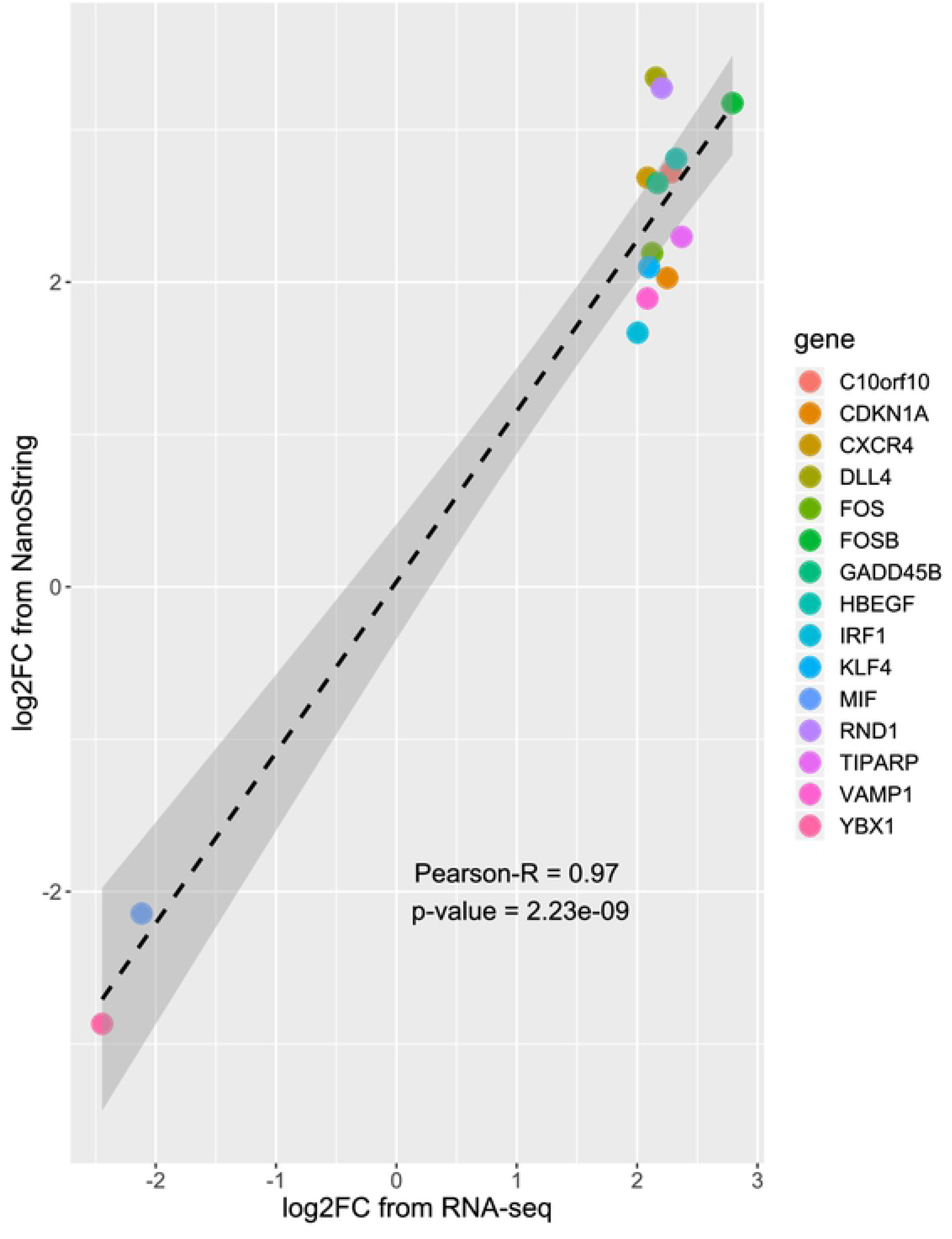
Comparison of RNA-seq and NanoString datasets. Both platforms displayed a strong correlation in gene expression (R^2^= 0.97; p=2.23E-09). FC: fold change.

## Discussion

Using a transcriptomic *in vitro* adenocarcinoma approach, we show that invasion of Caco-2 cells by pathogenic *F. nucleatum* strains results in the upregulation of host genes associated with tumorigenesis: *dll4* and *klf4*. Invasion by these isolates alone is enough to stimulate this response.

DLL4 is a Notch ligand that has been shown to play a role in angiogenesis and its expression facilitates tumour growth by accelerating vascular generation. This may provide insight into the mechanism of microbially-influenced tumorigenesis wherein the invasion of *Fusobacterium* serves to set event in motion that create a tumor permissive, proinflammatory microenvironment (reviewed by Xiu *et al.*(8)). Recent work has shown *dll4* expression in CRC tissue but not in adenomas(9). This result strengthens the association between the up-regulation of *dll4* upon *Fusobacterium* invasion and CRC development.

While *F. nucleatum* strain-induced upregulation of *klf4* was not as large as that for *dll4* when compared to *E. coli* infected control, its upregulation was significant compared to that observed for mock-treated cells. KLF4 is a transcription factor with known suppressive and oncogenic functions. How KLF4 functions is thought to be context dependent, though the mechanisms of this duality (be they posttranslational modifications, association with other proteins, subcellular location) are not understood (reviewed by Wang et al.(10)). Regardless, *klf4* has distinct functions in carcinogenesis as its upregulation has been linked to epithelial cell inflammation, cell proliferation and chemotherapy resistance (reviewed by Farrugia *et al.* 2016 (11)).

Given that *F. nucleatum* abundance has been directly correlated with susceptibility to chemotherapy failure and disease relapse (reviewed by Ternes *et al.*(3)), understanding potential mechanisms of microbially-associated tumorigenesis is of high importance for this bacterial species. The upregulation of both *dll4* and *klf4* highlights alternative mechanisms of action for this opportunistic pathogen to perpetuate CRC persistence and progression – possibly by regulation of angiogenesis, inflammation and cell proliferation.

## List of abbreviations

CRC: colorectal cancer
DEG: differentially expressed genes
*dll4*/DLL4: delta like canonical notch ligand 4
FC: fold change
*klf4*/KLF4: Kruppel-like factor 4
qPCR: quantitative polymerase chain reaction
RPKM: reads per kilobase of transcript, per million mapped reads

## Author involvement

KC and AVR: acquisition of data; analysis and interpretation of data; SDB: analysis of data; JP: drafting of manuscript and critical assessment; EA-V and RAH: obtained funding, study concept and design; study supervision.

## Transcript profiling

https://www.ncbi.nlm.nih.gov/geo/query/acc.cgi?acc=GSE130714

